# LeishMANIAdb: a comparative resource for *Leishmania* proteins

**DOI:** 10.1101/2023.03.08.531706

**Authors:** Gábor E. Tusnády, András Zeke, Zsófia E. Kálmán, Marie Fatoux, Sylvie Ricard-Blum, Toby J. Gibson, Laszlo Dobson

## Abstract

Leishmaniasis is a detrimental disease causing serious changes in quality of life and some forms lead to death. The disease is spread by the parasite *Leishmania* transmitted by sandfly vectors and their primary hosts are vertebrates including humans. The pathogen penetrates host cells and secretes proteins (the secretome) to repurpose cells for pathogen growth and to alter cell signaling via host-pathogen Protein-Protein Interactions (PPIs). Here we present LeishMANIAdb, a database specifically designed to investigate how *Leishmania* virulence factors may interfere with host proteins. Since the secretomes of different *Leishmania* species are only partially characterized, we collected various experimental evidence and used computational predictions to identify *Leishmania* secreted proteins to generate a user-friendly unified web resource allowing users to access all information available on experimental and predicted secretomes. In addition, we manually annotated host-pathogen interactions of 211 proteins, and the localization/function of 3764 transmembrane (TM) proteins of different *Leishmania* species. We also enriched all proteins with automatic structural and functional predictions that can provide new insights in the molecular mechanisms of infection. Our database, available at https://leishmaniadb.ttk.hu may provide novel insights into *Leishmania* host-pathogen interactions and help to identify new therapeutic targets for this neglected disease.

## Introduction

Leishmaniasis is a neglected tropical disease causing severe symptoms, affecting around 1 million new people yearly, with annual deaths estimated to be around 60,000 ***Torres et al. (2017)***. Although over 90% of cases occur in poor regions south of the Equator, due to climatic changes it also appears in new areas, and it has already shown up in Mediterranean European countries ***Gianchec***chi and Montomoli (***2020***) and Texas, USA ***McIlwee et al. (2018)***. To this date no approved human vaccine is available and treatment is most effective at an early stage of the infection. *Leishmania* parasites are unicellular, flagellated trypanosomatids, belonging to the class Kinetoplastea. Upon infection, the amastigote stage pathogen (with reduced flagella) is engulfed by phagocytes, where it ends up in a stable parasitophorous vacuole that protects it ***Arango Duque et al. (2019)***. *Leishmania* cells then proliferate unhindered within host cells until egress and spreading to nearby phagocytes ***Real et al. (2014)***. The parasite secretes proteins that enter various parts of the cell ***Atayde et al. (2015)***. The secreted virulence factors can then interfere with cell signaling by interacting with the host proteins: they increase glycolytic metabolism ***Ohms et al. (2021)***, perturb microbicidal pathways ***Matheoud et al. (2013)***, escape the innate immune response, and repurpose macrophages for parasite replication ***Atayde et al. (2016)*** by disturbing with cellular protein-protein interactions (PPIs). Interestingly, these mechanisms are somewhat unique to *Leishmania* among trypanosomes, which are usually extracellular pathogens and do not enter host cells. In contrast, *Leishmania* secretes proteins which are critical for host cell subjugation, but how they enter the cytoplasm of host cells is still poorly understood.

The host targeted interactions are often mediated via Short Linear Motifs (SLiMs) in many distant, unrelated intracellular pathogens, ranging from viruses and bacteria to unicellular eukaryotes ***Davey et al. (2011)***. SLiMs are flexible protein segments composed of a restricted number of residues (between 3-10), that usually bind to structured protein domains. Their short length and structural flexibility enable them to bind to a wide range of domains. Cellular SLiMs typically bind their targets with low micromolar affinity. These weak and transient interactions enable SLiMs to work in cooperative regulatory systems ***Van Roey et al. (2014)***. Pathogens mimic host SLiMs to interact with host cell proteins ***Davey et al. (2011)***. Pathogen SLiMs often bind with higher affinities than the cellular ones, outcompeting the native interactions, permanently re-wiring the host regulation network. A few modulatory SLiMs have already been discovered in eukaryotic pathogens, such as the Toxoplasma gondii MapK docking motif ***Pellegrini et al. (2017)*** and the stage-specific (promastigote-amastigote) phosphorylation motifs from *Leishmania* ***Tsigankov et al. (2013)***. In addition, several putative SLiMs were recently detected in *Leishmania* such as heparin-binding sequences or RGD integrin-binding motifs but their function has not been confirmed yet ***Peysselon*** et al. (***2013***).

Numerous studies investigated *Leishmania* secretomes. Most of them expose promastigotes to a heat shock and pH change (attempting to emulate the conditions that promote promastigote-to-amastigote stage transition) and then analyze the *Leishmania* conditioned medium by proteomics to identify secreted proteins ***Cuervo et al. (2009)***, and measure their protein abundance or by transcriptomics to detect mRNA levels ***Lahav et al. (2011)***. While high-throughput experiments inherently suffer from a certain level of noise, experiments on individual proteins may be more reliable in the case of *Leishmania* the vast majority focuses on leishmanolysin (GP63), a surface-anchored protease important for pathogenesis ***Gregory et al. (2008)***; ***Guay-Vincent et al. (2022)***. Furthermore, data were collected on different *Leishmania* species/strains identified via names and identiflers varying from one source to another, making a unified overview challenging. Another key step towards understanding the infection mechanism would be the identification of *Leishmania* surface proteins that can mediate the attachment of the pathogen to the host cell. Some surfaceome experiments were carried out on *Leishmania*-related species, and human host proteins binding to the surface of 24 strains of intact *Leishmania* have been identified ***Fatoux-Ardore et al. (2014)***. Beside the characterization of *Leishmania* secretomes, the identification of host-*Leishmania* PPIs is needed to narrow down virulence factors perturbing the host cell regulation to modules interfering with host proteins. SLiMs have low information content and simply scanning them in *Leishmania* secretomes may yield many false positives. Their structural and functional context, such as accessibility, conservation and localization, are all key elements to successfully identify those that may have a role in rewiring the host cell regulation. Notably, SLiMs also play a key role in maintaining housekeeping processes in *Leishmania*, therefore to find candidate SLiMs that may alter the host regulation, we need to discriminate SLiMs of proteins that reach the host cytoplasm or nucleus but limited information about these proteins are available. Currently the only publicly available database dealing with *Leishmania* proteins is TriTrypDB ***Shanmugasundram et al. (2023)***, which is part of the VEuPathDB ***Amos et al. (2022)***. TriTrypDB is a functional genomic resource for Trypanosomatidae, offering proteomic datasets, however it does not focus on protein structure, protein motif search and interactions.

We developed LeishMANIAdb to expedite *Leishmania* research by unifying scattered information from the literature in a user-friendly way and to extend available resources by adding protein level information. We collected high-throughput experiments and interaction studies on individual proteins, and used various prediction methods to enrich proteins with structural information.

## Results

### Selection of ***Leishmania*** proteomes and homology mapping of various kinetoplastid proteins

We selected 5 *Leishmania* species (reference proteomes: *L. brazliensis, L. donovani, L. infantum, L. major, L. mexicana*), 13 Leishmania strains (*Lbraziliensis MHOMBR75M2903, Lbraziliensis MHOMBR75M2904, Lbraziliensis MHOMBR75M29042019, Ldonovani BPK282A1, LdonovaniCL-SL, Ldonovani HU3, Ldonovani LV, Linfantum JPCM5, LmajorFriedlin, Lmajor Friedlin2021, Lmajor LV39c5, Lmajor SD75*.*1, Lmexicana MHOMGT2001U1103*), and 6 related species (reference proteomes: *Bodo saltans, Leptomonas seymouri, Trypansoma brucei, Trypansoma cruzi, Trypansoma rangeli, Trypanosoma theileri*) as an outgroup. The 5 *Leishmania* proteomes and the 6 related kinetoplastid proteomes were selected based on their quality (i.e. number of fragments and missing proteins) and were downloaded from UniProtKB ***UniProt Consortium*** (***2023***). *Leishmania* proteins were also cross-referenced to TriTrypDB ***Shanmugasundram et al. (2023)***. Around 30% of the cross-referenced proteins have different sequences deposited into these resources, and in most cases the difference is due to the position of the initiator methionine. For data compatibility we always use the UniProt sequence version but the conflicts are highlighted in LeishMANIAdb. We also performed a similarity search between these proteins and linked close homologs (see Methods) so annotations and predictions can be easily compared between them. All manual annotations and experimental data from different sources were mapped to these proteins. The 13 *Leishmania* strain proteomes were downloaded from TriTrypDB. Altogether LeishMANIAdb contains 40 537 searchable *Leishmania* proteins from reference proteomes, 108 766 proteins from different strains and 68 924 other kinetoplastid proteins to strengthen predictions.

### Manual annotation of host-pathogen PPIs and TM protein localization

We manually curated hundreds of proteins, using two strategies. The first type of annotation was the collection of host-pathogen PPI experiments on individual proteins, with the majority of them involving leishmanolysin (GP63). We collected 29 papers reporting 82 *Leishmania* PPIs with different hosts. Although experiments were mapped back to specific proteins, the results are also displayed on close homologs (with a note that the experimental data is derived from a different protein) resulting in 211 proteins that contain PPI data. Interactions were reported using the Minimum Information required for reporting a Molecular Interaction eXperiment MIMIx ***Orchard et al. (2007)*** community standard description. The second type of manually curated data was the localization and functional annotation of TM proteins. The aim was to find surface proteins that may facilitate the infection, but we annotated hundreds of other TM proteins with their localizations too. For this task, we used all close homologous proteins defined in the previous step. Altogether 342 protein families were annotated and these annotations were shared between 3764 proteins (which is 45.11% percent of the predicted TM proteomes, and 9.28% of all proteins of the 5 species combined).

### The definition of *Leishmania* secretome and protein localization is still incomplete

*Leishmania* not only exploits host-secretory pathways to distribute effectors but also utilizes an unusual mechanism to deliver proteins to the cytosol of infected cells by releasing exosomes into the parasitophorous vesicle, which might fuse with the vesicular membrane to release their protein content ***Silverman et al. (2010)***. Therefore computational methods based on signal peptides and localization predictions are not sufficient to predict *Leishmania* secretomes. To overcome this limitation we also used high-throughput experiments ***Silverman et al. (2008)***; ***Cuervo et al. (2009)***; ***Hassani et al. (2011)***; ***Forrest et al. (2020)***; ***Pissarra et al. (2022)*** to increase the coverage of *Leishmania* secretomes. Strikingly, the number of proteins in these secretomes varies to a large extent, and some proteins cannot be identified by mass spectrometry. Other datasets include proteins found in glycosomes ***Jamdhade et al. (2015)***, stage-dependent (promastigote/amastigote) phosphoproteomics ***Tsigankov et al. (2013)***, housekeeping gene localizations ***Jardim et al. (2018)***, exosome content ***Silverman et al. (2010)***, protein and mRNA abundance data ***Lahav et al. (2011)***; ***Pescher et al. (2011)***. When we mapped back all secretome and abundance experiments to *Leishmania infantum* (from orthologous proteins of other *Leishmania* species), the number of identified proteins ranges from 10 to 2,000 (Figure 1/A), and even when experimental conditions were similar they yielded highly different amounts of proteins. For example, pioneer secretome studies only provided a few hundred hits, while the latest ones are more inclusive with thousands of hits. Gene duplication is often acting on protein families responsible for host-pathogen PPIs, therefore we also collected proteins that are highly expanded. Notably, as all kinetoplastids have a polycistronic transcription system, the main way to amplify expression of critical proteins is through gene duplication. Thereby highly expanded gene families can be directly mapped to functions critical for these parasites ***Jackson et al. (2016)***. In this case we could discriminate between proteins with already many paralogs within kinetoplastids and *Leishmania*-exclusive amplified proteins. When we searched for homologs of *Leishmania infantum* proteins, we found distinct amino acid transporter and cofactor families already expanded in all kinetoplastids including *Leishmania*. In contrast, amastins, leishmanolysin, 3’A2-related proteins, kinase-containing putative receptor proteins (and several uncharacterized proteins) seemed to be highly abundant in *Leishmania* proteomes compared to all kinetoplastids (Figure 1/B). Comparing complete proteomic datasets yielded only a small overlap. We defined 1) *Leishmania*_novelty proteins, which are proteins without close homologs in SwissProt, without characterized Pfam domains, and expanded in *Leishmania infantum* (compared to other kinetoplastids); 2) abundant proteins, which are proteins showing increased abundance upon infection; 3) secreted proteins experimentally identified in at least two secretome experiments. These definitions provided markedly different protein sets, with some overlap between secreted and abundant proteins (611 proteins) and with only 22 proteins contained in all datasets (Figure 1/C).

**Figure 1.**
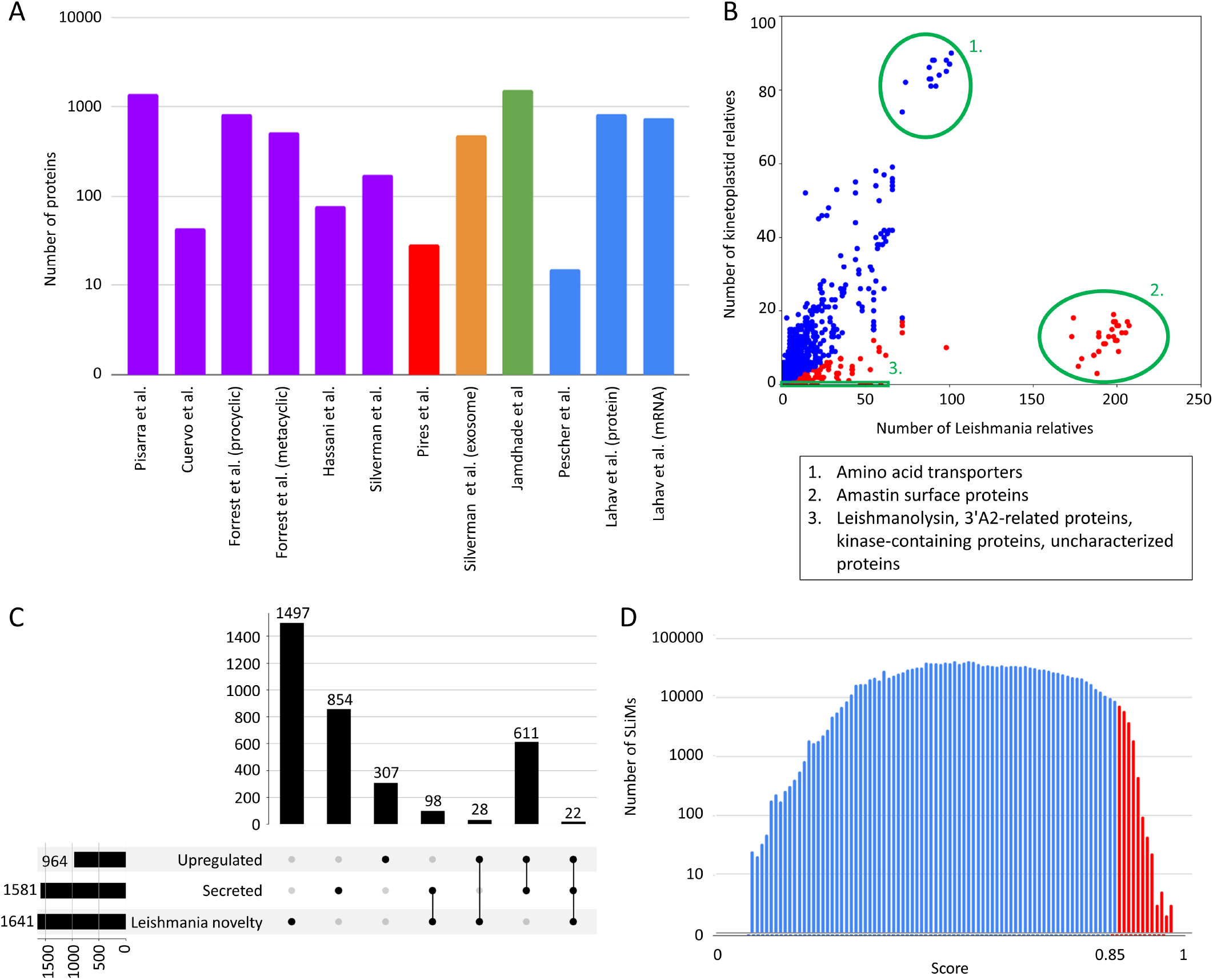
LeishMANIAdb content. All data were calculated on *Leishmania infantum*. A: Number of proteins in different proteomic datasets (purple: promastigote secretome, red: amastigote secretome, orange: exosome, green: housekeeping genes, blue: higher protein abundance level upon infection). B: Number of kinetoplastid and *Leishmania* close homologs. Each dot represents a protein (red: at least 80% of close homologs are in *Leishmania*, blue: other proteins). Green circles represent distinctive groups. C: Overlap between abundant, secreted and *Leishmania* novelty proteins (for more detail see text). D: Distribution of all predicted SLiMs with different scores. Red marks candidate motifs above 0.85 cutoff (for more details see text).

### AlphaFold2 provides an alternative way estimate structural features

We used different methods to predict the structural features of proteins. Classical sequence-based methods can detect globular domains ***Paysan-Lafosse et al. (2023)***, TM regions ***Dobson et al. (2015)*** and intrinsically disordered regions (IDRs) ***Erdős et al. (2021)***. However, the use of AlphaFold2 (AF2) ***Jumper et al. (2021)*** provides alternative ways to obtain structural information. In LeishMANIAdb we used structures available in the AlphaFold database ***Varadi et al. (2022b***) (however we could not find 3192 proteins (6% of all *Leishmania* proteins in LeishMANIAdb)). We not only displayed the predicted 3D structure of the proteins, but also information derived from the AF2 models such as the secondary structures and the position of the lipid bilayer for membrane proteins using the method introduced in the TMAlphaFold database ***Dobson et al. (2023)***. Although AF2 was originally built to predict protein structure, the scientific community quickly realized it is as much (if not more) efficient at predicting protein disorder ***Akdel et al. (2022)***. To analyze IDRs we displayed predicted local distance difference test (pLDDT) values and relative surface accessibility from AF2. For IDR prediction in TM proteins, we tailored MemDis ***Dobson and Tusnády*** (***2021***) to incorporate features from AF2 instead of sequence-based predictors (see Methods).

### Short linear motif candidates that may hijack host cell regulation

We scanned *Leishmania* proteins for SLiMs using the regular expressions stored in the Eukaryotic Linear Motif (ELM) resource ***Kumar et al. (2022)***. Scanning SLiMs alone would mostly yield false positive hits, so we developed a scoring system that ranges from 0 to 1, and that takes into account most information we collected. We aimed to develop a scoring system where conservation and accessibility/disorder has a reasonably high weight, while keeping in mind that proteomic experiments and localization information are a good way to narrow down the potentially large number of false positive hits. Unfortunately, due to the lack of data, in the case of *Leishmania* it is not possible to construct a benchmark set to evaluate motif scores. We can still assume that a good starting point can be when most predictions and proteomic data agree. Considering *Leishmania infantum* alone, we detected over a million putative motifs, from which 1.21% had a score above 0.85, on 343 proteins (Figure 1/D).

### The LeishMANIAdb web resource

To visualize all the collected and calculated information we developed an open-access resource. In LeishMANIAdb users can search for proteins using their UniProt Accession (AC), Entry name (formerly ID), gene name and protein name. We also provide several protein sets as examples to help users browsing the database. Currently proteins are sorted based on 1) species: L. braziliensis, L. donovani, L. infantum, L. major, and L. mexicana; 2) manual curation data; 3) experimental data: secreted proteins, protein abundance/mRNA level data, proteins with any kind of experimental data listed above; 4) computationally predicted information: proteins expanded in *Leishmania* (score >= 0.8 see Methods, Supplementary Material), transmembrane proteins, proteins with high disordered content (at least 70% predicted disorder), proteins with high-scoring linear motifs (score >= 0.85) and novel kinetoplastid proteins (proteins without SwissProt close homologs or Pfam domains). After searching (or selecting a protein set) users can further narrow their selection by choosing any other criterion (Figure 2/A).

**Figure 2.**
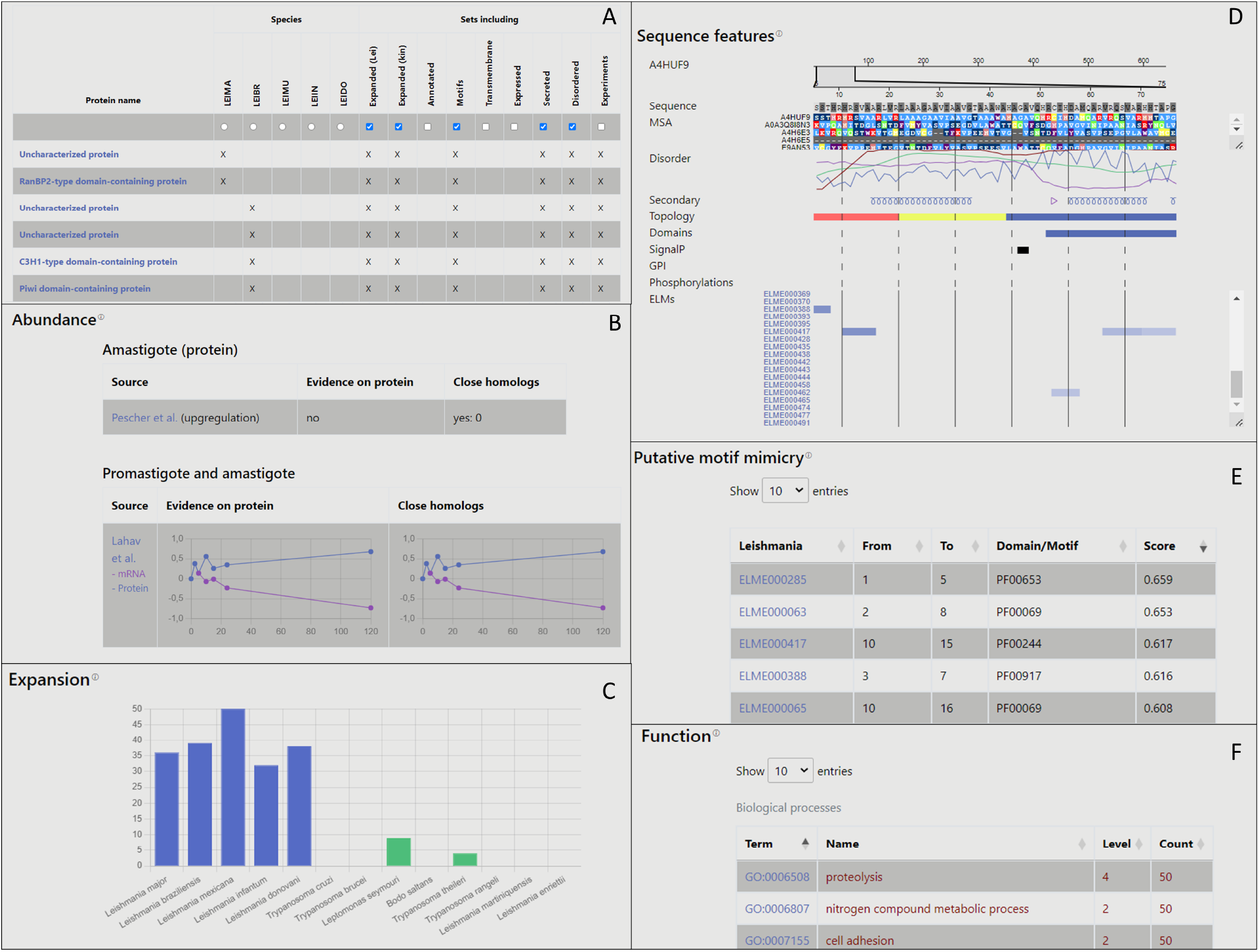
Figure 2: Layout of LeishMANIAdb: A: the search/browse result menu. B: Expression section of the entry page. C: Expansion section. D: Sequence features section. E: Putative motif mimicry section. F: Function section.

The entry page for proteins consists of 10 sections, which are only visible if they contain data. The “Quick info” displays the protein name, species, cross-references and its number of amino acid residues. Data curation appears under the “Annotations” section. PPI (curated at the MIMIx level), localization and function annotations are mirrored from close homologous proteins. We also display functional annotations by Jardim et al. ***Jardim et al. (2018)***. The “Localization” section contains high-throughput experiment data - promastigote and amastigote secretion, an exosome experiment and the glycosome. Protein localization, signal peptide and glycosylphosphatidyl (GPI) anchor predictions are also displayed here. Since the reliability of both predictions and experiments may vary, we also display all this data for close homologous proteins, so users can quickly check the robustness of information. Furthermore, we also collected Gene_Ontology (GO) ***The Gene On-*** tology Consortium (***2019***) annotations for cellular compartments. In this case, the specificity of the term (how deep it is on the tree) is shown in the level column. GO annotations are collected for all close homologous proteins too, and the number of occurrences of each term is displayed. We highlighted terms that are associated with the inspected protein itself that is displayed on the page. The “Abundance” module can display the mRNA and protein level experiments: static/single point (upregulated or not) or time-course experiments (e.g., mRNA and protein levels available for 7 timesteps across 120 hours, Figure 2/B). In the “Expansion” section the number of close homologs are displayed by species with a color-code to identify *Leishmania* intracellular and extracellular/freeliving relatives (Figure 2/C). The “Sequence features” displays various information (Figure 2/D). On the top, the gapless multiple sequence alignment (MSA) of proteins from the reference proteomes is visible. In this alignment gaps from the entry protein were removed (the original alignment with all strains can be downloaded) so other protein features could be visualized. Protein disorder, secondary structures, transmembrane topology prediction, domains, signal peptides and GPI anchors and stage-dependent phosphorylation are also displayed. Predicted SLiMs are shown with a colorcoded score (see Methods, Supplementary Material). In the “Structure” section the AF2 predicted structure is available (with the position of the membrane domain for TM proteins). The “Putative motif mimicry” section is the table format version of SLiMs from the “Sequence features” module (Figure 2/E). The “Function” section contains GO Molecular Function and Biological Process terms.

As done for the Cellular Component, terms are mirrored from close homologous proteins, and they can be sorted based on their specificity (how deep they are located in the tree) and occurrences considering homologs (Figure 2/F). Finally, each result from a BLAST search against SwissProt and kinetoplastids relatives are listed in the “Homologs” table.

For each protein the full MSA, high-throughput experiments, annotation and predicted sequential features can be downloaded from the bottom of the page. Batch download is also available to download the full database or different protein sets.

## Discussion

### Reliability of data

In LeishMANIAdb we aimed to collect high-throughput experimental data, PPI data on individual proteins, predictions and localization information based on distant homologs. We noticed that the amount of data from MS experiments differs highly, and therefore likely the quality also varies. Not unexpected from high-throughput techniques applied to less-studied organisms, this can be attributed to the quality of sample preparation, MS experimental techniques and most likely to the sequences in the background databases. One striking finding was that the secretory datasets contain a large number of proteins that are likely to take part in the housekeeping processes of *Leishmania* cells, such as cytoskeletal proteins, nuclear histones and metabolic enzymes. Exosomes are known to contain a relatively high amount of "background” proteins leaking from the cytosol of cells. Another explanation is that several housekeeping genes (such as intracellular chaperones and enzymes) are moonlighting proteins, they are generally constitutively expressed and have high levels of expression, while they are fulfilling other functions outside the cells ***Jeffery*** (***2018***). Due to the lack of comparative studies, we cannot assess the enrichment ratios of secreted molecules, to see if there is selective exosomal packaging of a well-defined subset of *Leishmania* proteins. However, *Leishmania* exosomal-like secretion also differs from the typical exosomal sorting seen in other eukaryotic organisms because budding primarily initiates at the cell membrane, and not inside multivesicular bodies (endosomes). Therefore it is equally possible that in Leishmaniids, the budding is non-selective for its cytoplasmic cargos. Instead it would be initiated by cell surface receptors and primarily serve as a defense mechanism against membrane-attached host complement and other immune complexes, removing them before they could damage the parasite membrane. Currently, testing of the latter hypothesis is impossible, since only soluble components but not the integral membrane proteins of *Leishmania* exosomes have been studied in depth in the above cited studies.

From a computational point of view, predicting any features on *Leishmania* proteins might be highly challenging, as methods established were mostly trained on sequences that show little or no similarity to *Leishmania* proteins. The 5 *Leishmania* reference proteomes contain 10 267 uncharacterized proteins combined, which is 25% of LeishMANIAdb. TMAlphaFold provides an objective quality measurement option for alpha-helical membrane proteins. When we compared the TM proteome of Homo sapiens and *Leishmania infantum* we noticed that the ratio of good and excellent quality structures was much lower in *Leishmania*, probably caused by the different coverage of kinetoplastid and human structures deposited into the PDB (Figure 3/A).

**Figure 3.**
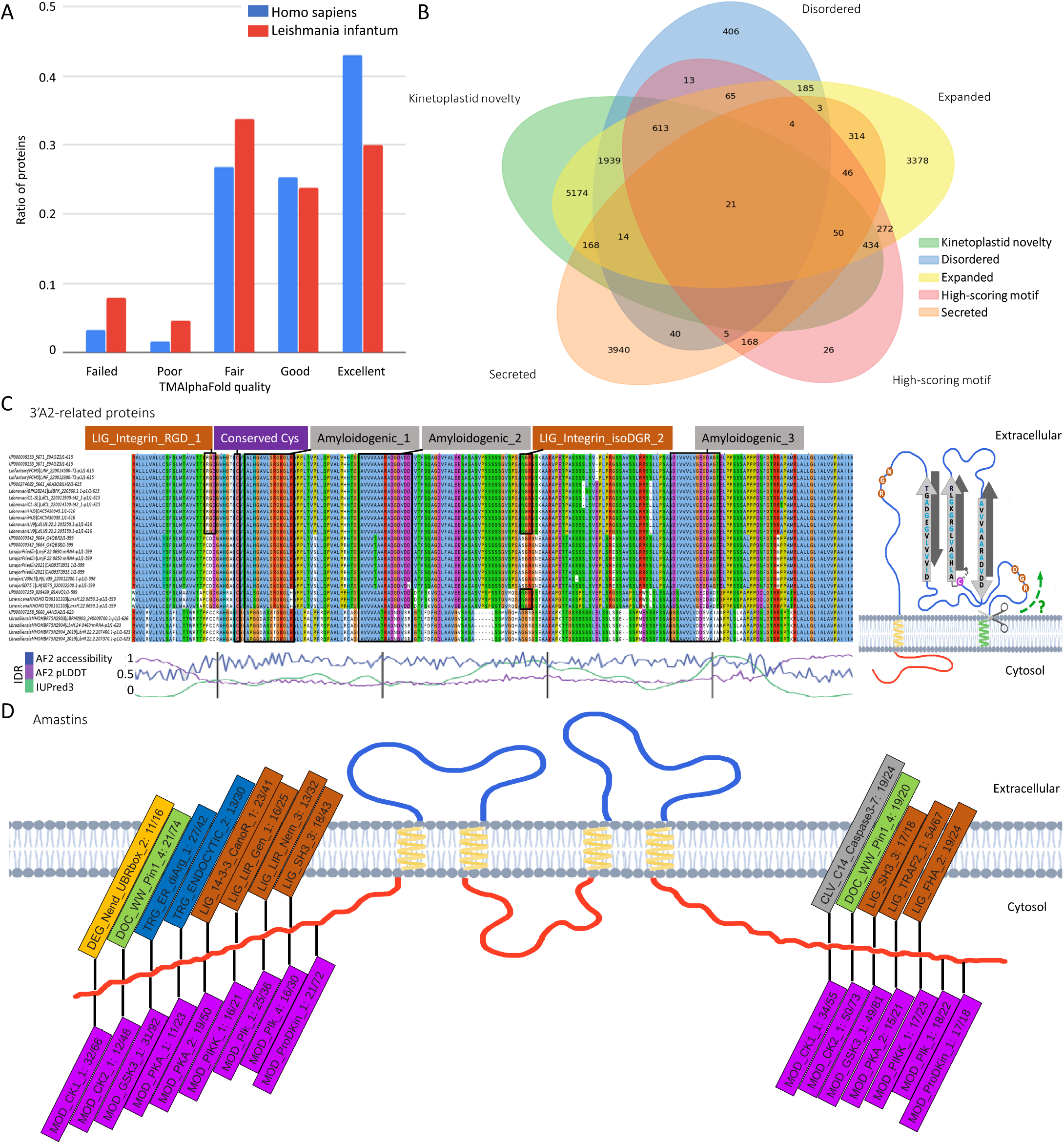
Figure 3: A: Distribution of membrane protein quality levels of AlphaFold structure in Homo sapiens and *Leishmania infantum*. B: Venn diagram of proteins that are 1) secreted 2) novel kinetoplastid 3) expanded (or new) in *Leishmania* 4) disordered 5) contain candidate SLiMs. C: left: Multiple Sequence Alignment of 3’A2 related proteins (alignment is available under UniProt AC: E9AGZ3). Amyloidogenic regions, conserved cysteine and Integrin-binding motifs are highlighted; right: proposed topology of 3’A2 related proteins D: Frequent SLiMs in the cytoplasmic tail regions of amastins (the numbers denote the unique/total occurrences).

### Case studies

LeishMANIAdb can be utilized for different purposes, and can be a good starting point for various analyses. We selected three examples that highlight some use cases of the resource.

Using the Browse menu, after selecting a category users can further narrow down their search for proteins selecting additional categories to refine the results. For instance, if users are looking for *Leishmania* SLiMs that may alter or rewire host cell regulation network, they can look for proteins that were experimentally proven to be secreted, and then select proteins with disordered regions because SLiMs are mostly located in IDRs. “Kinetoplastid novelty” selection ensures that the protein and its domains are not present in other organisms, while *Leishmania* novelty/expansions select proteins that are new or highly expanded in *Leishmania* species. Last, by selecting highscoring motifs users get a list of proteins where the motif is most likely to be functional (Figure 3/B shows the Venn diagram of the selection). These proteins may be an interesting starting point for further analyses.

When performing systematic searches to identify possible parasite hits of integrin ligand motifs (that only functions in the host, as kinetoplastids have no integrins), we identified a striking set of examples in a family of poorly-known *Leishmania* genes called 3’A2 related ORFs. This kinetoplastidspecific family of genes is actually expanded in *Leishmania* species together with the canonically secreted A2 proteins, which are known pathogenicity factors ***Zhang and Matlashewski*** (***2001***). While the actual sequences of these proteins are poorly conserved and very little is known about their subcellular location, the *Leishmania* versions have at least one transmembrane region and a Cterminal cytoplasmic tail, with an N-terminal signal peptide (or possibly another TM segment). Nevertheless, in the predicted, largely disordered extracellular segment we observed multiple, short, conserved stretches that may have amyloidogenic properties (high Val, Ala and Gly content, upon visual inspection), presumably capable of oligomerization and amphiphilic interaction with membranes (Figure 3/C). A highly conserved cysteine residue preceding the first amyloidogenic sequence might help the homodimerization by forming a disulphide bridge with neighboring 3’A2 related protein. Strikingly, in *Leishmania infantum* and *Leishmania donovani* (both species capable of causing visceral leishmaniasis), the N-terminus of these proteins carries canonical RGD (Arg-GlyAsp) sequences, immediately after the putative signal peptide cleavage site. In addition, *Leishmania donovani* and *Leishmania infantum* proteins contain an NGR motif where asparagine deamidation might yield an isoDGR motif. If these proteins are expressed on the cell surface, they might bind to host integrins in an oligomeric state, and might even attack the host membrane as if it were a beta-barrel pore-forming toxin. However, much more experiments are needed to test any of these hypotheses.

Amastins are a large family of kinetoplastid-specific membrane proteins that belong to the broader claudin-like superfamily, implicated in the maintenance of parasitophorous vacuoles ***de*** Paiva et al. (***2015***). Accordingly, the majority of amastins have 4 tightly packed TM segments, with cytosolic tail regions. Similarly to their vertebrate counterparts that form tight cell-cell junctions by complex oligomerization processes, amastins might also engage in a variety of interactions with internal as well as external, host proteins. Although their exact function is not known, among the 221 identified amastins with 4 TM regions we looked for SLiMs that occur in multiple proteins. Predicted SLiMs (within disordered regions) were packed in their cytoplasmic tail regions (Figure 3/D). Since these regions face inward the parasite, we further narrowed hits based on their binding domain to be present in *Leishmania*. We identified multiple potential phosphorylation sites and protein-protein interaction motifs, such as SH3 ligands (*Leishmania* species do encode SH3 domain proteins) as well as vesicular trafficking signals. The tail region of amastins seem to be highly variable, likely acting as a hotspot in the pathogen-host arms race.

### Comparison with other resources

In the past decades several databases were built to investigate *Leishmania*, however they are unfortunately often offline and no longer updated by now. LeishCyc ***Saunders et al. (2012)*** focused on biochemical pathways. LeishDB ***Torres et al. (2017)*** included coding genes and non-RNAs and provided new annotation to them. The cysteine protease database in *Leishmania* species ***Rana*** et al. (***2012***) was designed to find data related to cysteine protease and LeishBase was a structural database. There are a few active databases: Leish-ExP (http://www.hpppi.iicb.res.in/Leishex) (which has not so far been published in a peer-reviewed journal) contains proteins exclusively present in *Leishmania*. Leish-ExP incorporates localization tools, includes GO annotations and calculates physico-chemical properties of proteins. LmSmdB ***Patel et al. (2016)*** focuses on metabolic and biosynthetic pathways. TriTrypDB ***Shanmugasundram et al. (2023)*** is a kinetoplastid database that is part of the VEuPathDB resource ***Amos et al. (2022)***. These databases contain a lot of experimental data and various tools to analyze eukaryotic pathogens, but they are mostly focused on genomic data although proteomic datasets, and some protein prediction algorithms are also incorporated.

There are also a handful of databases that include information on host-pathogen interactions: HPIDB ***Ammari et al. (2016)***, PHIDIAS ***Xiang et al. (2007)*** and PHI-base ***Urban et al. (2022)*** contain information about PPIs between the host and pathogen, while ImitateDB ***Tayal et al. (2022)*** specifically focuses on motif mimicry. These resources contain no or very little data about *Leishmania*.

In LeishMANIAdb our main goal was to include protein information relevant to the infection and to complement previously established and still available resources. We included several proteomic datasets, and enriched experimental information with state-of-the-art prediction tools. Still, the most powerful way to explore uncharted proteomes is to inspect MSAs and check for conserved residues and regions - LeishMANIAdb contains precalculated alignments for all proteins. We also added hundreds of annotations to thousands of proteins, including localization and interaction information. While several databases seem to be shut down after a couple of years, our laboratory hosts several resources and we routinely update them. We plan to do so with LeishMANIAdb as well as to expand its repertoire to host-*Leishmania* interactions involving glycans and glycolipids, which play major roles in the infection.

## Methods

### Resources

Protein sequences were retrieved from UniProtKB (release 2022_05) ***UniProt Consortium*** (***2023***) and from TriTrypDB ***Shanmugasundram et al. (2023)*** based on the UniProt cross-references (*L. braziliensis, L. donovani, L. infantum, L. major, L. mexicana, Bodo saltans, Leptomonas seymouri, Trypansoma brucei, Trypansoma cruzi, Trypansoma rangeli, Trypanosoma theileri*). Homologs in other kinetoplastids and in SwissProt were searched with BLAST using e-value: 10-5; sequence identity>20%; coverage>50%. In the “Homologs” section all results are displayed, however for most other sections (and calculation) we only used homologous proteins until the first non-kinetoplastid SwissProt hit considering sequence identity (termed as “close-homologs”); Further similar kinetoplastid proteins were therefore considered as a different homology group. Furthermore, we downloaded strains belonging to the 5 selected *Leishmania* species from TriTrypDB. In this case a more stringent condition was used in BLAST, by setting E-value: 10-5; sequence identity>80%; coverage>80%. All kinetoplastid species and strains were used to calculate motif conservation.

We prepared three different type of MSAs using ClustalOmega ***Sievers et al. (2011)***: 1) “nonredundant” MSA using homologous proteins from kinetoplastid reference proteomes; 2) the same MSA but with gaps removed from the “reference” protein that is currently displayed on the webpage; 3) a more redundant MSA using homologous kinetoplastid proteins in all species and strains (used to calculate motif conservation).

High-throughput experiments were first mapped to the corresponding protein using the identifler provided in the original paper, then mirrored to close *Leishmania* homologs if their sequence were identical.

IDRs were predicted using IUPred3 ***Erdős et al. (2021)*** and using the AF2 models’ pLDDT and accessibility values - the latter was calculated by DSSP ***Joosten et al. (2011)***, normalized using maximum values calculated as in Tien et al. ***Tien et al. (2013)***, the exposed value threshold defined as suggested by Rost et al. ***Rost and Sander*** (***1994***). In the case of TM proteins, IDRs were also predicted by MemDis ***Dobson and Tusnády*** (***2021***). In this in-house modified version, the Position-Specific Scoring Matrices (PSSMs) were generated using kinetoplastid sequence library and secondary structure and accessibility were calculated using AlphaFold2. Topology was predicted by CCTOP ***Dobson et al. (2015)***, however to minimise sporadic erroneous predictions, after an initial prediction we performed a constrained iteration where the topologies of homologous proteins were used as a constraint. Using this approach, closely related proteins will likely have the same topology. Secondary structure elements derived from AF2 structures are also displayed. Pfam domains were identified using InterPro ***Paysan-Lafosse et al. (2023)***. Protein localization was assigned by GO ***The Gene Ontology Consortium*** (***2019***), predicted by DeepLoc ***Almagro Armenteros*** et al. (***2017***) and SignalP6.0 ***Teufel et al. (2022)***. NetGPI ***Gíslason et al. (2021)*** was used to predict GPI-anchors (all prediction results are visible, therefore in case of a contradiction it is up to the user to judge the results).

To detect SLiMs that may alter or rewire host cell regulation, we used the regular expressions from ELM ***Kumar et al. (2022)*** on all *Leishmania* sequences. We defined different context fllters and merged them into a single score to rank motifs (for more details see Supplementary Material): 1) Disordered: The score is the average of the IUPred3, AF2-based pLDDT and accessibility values. These disordered scores were first transformed so they range from 0 to 1, with 0.5 being the threshold, before calculating their mean; 2) Conservation of the motif was checked among close homologs with some permission for slight misalignment, and penalizing motifs that are present across all kinetoplastids - notably, in this case proteins from different *Leishmania* strains were also considered; 3) Localization: we used a simplified (intracellular/extracellular) distinction. Motif localization was determined using ELM GO annotations, secretion information and CCTOP prediction, while the domain localization was determined from TOPDOM (Varga et al., 2016). We looked for motif-domain pairs where they both have the same simplified (in/out) localization; 4) mRNA level: using transcriptomic experiments about expression data; 5) protein level: from experiments about protein abundance; 6) Secretion score based on secretome experiments; 7) Expansion score: reflecting how much the protein is expanded in *Leishmania* species (strains not included) compared to all kinetoplastids; 8) Outgroups score favoring proteins without homologs in SwissProt. Structure data reflects structure data deposited in the PDB ***Varadi et al. (2022a***) before 26.03.2023 and the AlphaFold database (v3). All other data was downloaded in October, 2022 from the source databases.

### Manual curation

We manually curated hundreds of proteins, using two strategies. First, we searched PubMed and Google scholar for “*Leishmania* host-pathogen protein interaction” and manually processed the results. Each protein in the experiments was mapped to the corresponding UniProt entry. Then we mapped interaction data to the 5 *Leishmania* proteomes. When the experiment was performed on a protein from different species, we mirrored it to the closest homology group in LeishMANIAdb, and we also indicated on the webpage that the experiment is from a different protein. All interactions were reported according to the community standard MIMIx level ***Orchard et al. (2007)***.

Next we searched for possible surface proteins. For this task we considered all homologs. We collected topology prediction, GPI-anchor prediction, GO terms and also searched for homologous proteins that were measured ***Bausch-Fluck et al. (2018)***; ***Langó et al. (2017)*** or predicted to be on the surface. We manually processed the entries using this approach, taking distant homologues, domain architectures and conservation patterns into consideration.

### Website design

The LeishMANIAdb website is written in PHP (v8.0) using the Laravel (v9.19) framework. All downloaded, predicted or calculated data are stored in a local MySQL (v8.0) database. To visualize sequence features over amino acid sequences, we developed a javascript package using React (18.2), while 3D structures are visualized using the original (for non-TM proteins) or a locally modified version of Mol* ***Sehnal et al. (2021)*** for TM proteins. The modified version can display the membrane as two planes around the investigated TM protein using the results of TMDET ***Tusnády et al. (2005)***.

## Acknowledgments

This project has received funding from the European Union’s Horizon 2020 research and innovation programme under the Marie Sklodowska-Curie grant agreement No 101028908; Ministry of Innovation and Technology of Hungary from the National Research, Development and Innovation Fund [K132522]. The authors would like to thank the help of the Gibson team at the European Molecular Biology Laboratory, especially Jesus Alvarado-Valverde for helpful suggestions and discussions, and Philippe Esterre (Pasteur Network, Institut Pasteur, Paris) for his help in selecting a number of *Leishmania* interactions to be curated.

