## Supplementary Material for "LeishMANIAdb: a comparative resource for *Leishmania* proteins" (from another version of this preprint: version 2, 2023-04-19)

### Scoring motifs

Residue-based scores

1. Disordered score

Disordered score was calculated using the following formula

$$Motif_{disordered_{score}} = \frac{\sum_{i=start}^{end} method\_score_i}{n(end-start)}$$

Where $n = number\ of\ methods$; *start, end* marks the starting and ending position in the motif, respectively. In most cases three methods were incorporated (sometimes AlphaFold2 structure was missing).

$$IUPred = output\ probability\ of\ IUPred\ for\ a\ given\ position$$

$$AlphaFold2_{acc} = \{\frac{RSA}{0.36*2},\ if RSA > 0.36\ 0.5 + \frac{(RSA+0.36)}{(1-0.36)*2},\ else$$

Where Relative Surface Accessibility (RSA) was calculated by normalizing DSSP accessibility output using values determined by Tien et al. The 0.36 threshold for exposed residues was determined by Rost et al. The equation yields 0.5 score at 0.36 value, and intermediate values between 0-0.36 and 0.36-1 are evenly rescaled.

$$AlphaFold2_{pLDDT} = \{\frac{\left(1-\frac{pLDDT}{100}\right)}{(1-0.7)*2},\ if\ pLDDT > 70\ 0.5 + \frac{\left(1-\frac{pLDDT}{100}\right)-(1-0.7)}{0.7*2},\ else$$

Where pLDDT is the confidence value for a given residue from AlphaFold2. The equation yields 0.5 score at 70 value, and intermediate values between 0-70 and 70 -100 are evenly rescaled. Note, that pLDDT inversely correlates with protein disorder.

Furthermore, any region listed in DisProt automatically gets maximum score (1). Transmembrane regions, signal peptides and PFAM domains get minimum score (0).

2. Conservation score

Conservation score was calculated on aligned Short Linear Motif pairs, when their distance were below 20 residues (permitting alignment errors)

$$\frac{\sum_{i=aligned\_motifs\_leishmania} 1-\frac{distance_i}{20}}{\sum_{i=aligned\_motifs\_trypanosomas} 1-\frac{distance_i}{20}}$$

Where $distance_i$ is the distance between two aligned motifs. The score is the highest when a motif is present in Leishmania, but missing from other Trypanosomas.

3. *Localization score

Localization score can be only calculated when there is evidence for both the motif and the corresponding domain localization. Currently there is a two-stage version was applied, where motifs and domains can be intracellular and extracellular.

Motif localization was determined using ELM GO annotations, Protein secretion information (high-throughput experiments and SignalP), and CCTOP prediction (in case of contradicting information this order was followed). Domain localization was determined using TOPDOM.

If the localization on the domain and motif side are the same, the score is 1, otherwise 0. (if there is not enough information this metric was not taken into account)

Protein scores:

4. *Maximum normalized mRNA expression score (using experiments by Lahav et al.)
5. *Maximum normalized protein expression score (using experiments by Lahav et al.)
6. $Secretion\ score = \frac{\sum_{i=sets\_for\_species} x_i}{n}$

   Where n is the number of sets covering the species and

$$\{x_i = 1,\ if\ included\ in\ the\ secretion\ set\ x_i = 0,\ if\ not\ included\ in\ the\ secretion\ set$$

7. $Expansion\ score = \frac{\sum_{i=leishmania\_species} x_i}{\sum_{i=trypanosoma\ species} x_i}$

   Where

$$\{x_i = 1,\ if\ protein\ present\ in\ the\ species\ x_i = 0,\ else$$

8. Outgroup score: Penalizing proteins with homologs in non-Trypanosomes. For this step all proteins were searched agains SwissProt using BLAST.

$$outgroup\_score = 1 - highest\_sequence\_identity$$

Scores with * cannot be applied for all proteins or motifs. The weight of different scores is depending on the scores that can be calculated (see tables below).

Residue-based scores (total weight 0.7)

| | With localization score | No localizations score |
|---|---|---|
| Disordered score | 0.25 | 0.35 |
| Conservation score | 0.25 | 0.35 |
| Localization score | 0.2 | 0 |

Protein scores (total weight: 0.3) :

| | With expression scores | No expression scores |
|---|---|---|
| mRNA expression score | 0.06 | 0 |
| Protein expression score | 0.06 | 0 |
| Secretion score | 0.06 | 0.1 |
| Expansion score | 0.06 | 0.1 |
| Closest homolog | 0.06 | 0.1 |
